# Achilles Tendon Structure Differs Between Runners And Non-Runners Despite No Clinical Signs Or Symptoms Of Mid-Substance Tendinopathy

**DOI:** 10.1101/290866

**Authors:** Todd J. Hullfish, Kenton L. Hagan, Ellen Casey, Josh R. Baxter

## Abstract

Achilles tendinopathy affects many running athletes and often leads to chronic pain and functional deficits. While changes in tendon structure have been linked with tendinopathy, the effects of distance running on tendon structure is not well understood. Therefore, the purpose of this study was to characterize structural differences in the Achilles tendons in healthy young adults and competitive distance runners using quantitative ultrasound analyses. We hypothesized that competitive distance runners with no clinical signs or symptoms of tendinopathy would have quantitative signs of tendon damage, characterized by decreased collagen alignment and echogenicity, in addition to previous reports of thicker tendons. Longitudinal ultrasound images of the right Achilles tendon mid-substance were acquired in competitive distance runners and recreationally-active adults. Collagen organization, mean echogenicity, and tendon thickness were quantified using image processing techniques. Clinical assessments confirmed that runners had no signs or symptoms of tendinopathy and controls were only included if they had no history of Achilles tendon pain or injuries. Runner tendons were 40% less organized, 48% thicker, and 41% less echogenic compared to the control tendons (p < 0.001). Young adults engaged in competitive distance-running have structurally different tendons than recreationally-active young adults. While these structural differences have been associated with tendon damage, the lack of clinical symptoms of tendinopathy may suggest that these detected differences may either be precursors of tendinopathy development or protective adaptations to cyclic tendon loading experienced during running.

## Introduction

Mid-substance Achilles tendinopathy is among the most commonly reported injuries in running athletes [1]. Repetitive tendon loading as high as 12 times body weight [2] is thought to elicit structural changes observed in tendinopathy development. This overuse injury presents with pain and swelling of the tendon4-6 cm proximal to the calcaneal insertion, which leads to upwards of a 50% decrease in tendon material properties [3]. Qualitative ultrasonography is often used to confirm this diagnosis using qualitative grading scales [4–6]. Once symptomatic, Achilles tendinopathy can be challenging to treat and may result in to sub-optimal clinical outcomes [7]; therefore, it is essential to develop sensitive predictors of pre-symptomatic tendinopathy. While structural changes, in both human and animal tendon, have been characterized using histological and imaging techniques [8], these invasive measurements are not practical to employ in clinical settings.

*In vivo* measurements of tendon structure have been linked to tendon damage in both human and small animal studies. Magnetic resonance imaging has highlighted tendon hypertrophy in response to tendon loading during running in elite and recreational athletes [9,10]; however, these observations have only characterized the cross-sectional area of the tendon – not the underlying structure. Micro-morphological analyses of symptomatic Achilles tendons are correlated with decreased tendon mechanics [11] but have not quantified structural differences in individuals at increased risks of developing tendinopathy. Crossed polarizer imaging [12,13] and an ultrasonography analogue [13–15] detect structural changes, characterized by decreased collagen alignment, in response to acute tendon damage and the healing response in small animal models. However, the order of structural changes and symptomatic tendinopathy have not yet been defined. The first step to elucidating this structure-symptom relationship is to quantify tendon structure in a cohort of individuals that are at an elevated risk of developing tendinopathic symptoms [16].

The purpose of this study was to quantify Achilles tendons structure in recreationally active adults and competitive distance runners using a repeatable and reliable ultrasound measurement of collagen organization [accepted manuscript]. We hypothesized that competitive distance runners will show quantitative signs of tendon damage, characterized by decreased echogenicity [17] and collagen alignment [12,13], and have thicker Achilles tendons [10] compared to recreationally active adults, despite no reports or clinical signs of tendinopathy symptoms. A secondary aim of this study was to determine if sex, body mass index (BMI), foot strike pattern, or years running affect collagen organization and echogenicity in competitive distance runners.

## Methods

Twenty-two competitive-collegiate distance runners (12 males/10 females; Age: 19 ± 1.5, BMI: 20.3 ± 1.6) and twelve healthy controls (5 males/7 females; Age: 25 ± 2; BMI: 23.8 ± 2.4) participated in this IRB approved study (Table 1). All subjects met several inclusion criteria: no Achilles tendon pain at or three months prior to the initial evaluation, no history of partial or complete tendon rupture, and ability to perform single leg hops on both sides. Age, sex, height, and weight were collected from all subjects. In addition to demographics, other information was collected from the runner group: a clinical outcome questionnaire (VISA-A), self-reported running foot-strike pattern, lower-extremity injury history, years participated in collegiate cross country and/or track and field. All competitive-distance runners participated on an NCAA cross-country team and images were acquired during a single session prior to the start of the competitive season. Individuals with clinical signs of tendinopathy identified through qualitative ultrasonography assessments and self-reported clinical outcomes and excluded from analysis. Longitudinal B-mode ultrasound images of the right Achilles tendon were acquired while subjects lay prone on a treatment table with the anterior aspect of the shank resting on the table and feet hanging off the table in the resting position. These images were acquired at the mid-substance of the Achilles tendon, approximately 4 cm proximal to the calcaneal insertion, using an 18 MHz transducer (L18-10L30H-4, SmartUs, TELEMED, Vilnius, Lithuania) with a scanning width of 3 cm (scan parameters: Dynamic Range: 72dB; gain: 47 dB). Longitudinal Doppler images were acquired in the runner group as part of the clinical evaluation to quantify the level of neovascularization of the tendon [6]. All images were acquired by a single investigator, saved as videos, and evaluated by a fellowship-trained sports medicine physician who was blinded to all other subject data. Tendon structure and neovascularization were both graded [6] on a scale from 0 to 3: ranging from normal (0) to severe symptoms (3). Recreationally active adults served as the control group in this study and reported no history of Achilles tendon pain or injuries and therefore were not evaluated using a clinical grading system [6].

**Table 1.** Subject demographics

| | **Sex**<br>Female/Male<br>(total) | **Age**<br>Mean ± STD<br>(years) | **Height**<br>Mean ± STD<br>(cm) | **Weight**<br>Mean ± STD<br>(kg) |
| --- | --- | --- | --- | --- |
| Runners | 9/10 (19) | 19 ± 1.5 | 172 ± 7 | 60.4 ± 8 |
| Controls | 7/5 (12) | 25 ± 2 | 175 ± 7 | 73 ± 9 |

Collagen organization was quantified in the longitudinal B-mode ultrasound images using custom-written software that has been described in detail elsewhere [13]. Briefly, a computational analogue for crossed polarizer imaging was employed to quantify collagen fascicles alignment of the mid-substance of the Achilles tendon. Fascicles are hyperechoic compared to the hypoechoic extracellular matrix which gives tendon a banded appearance. The orientations of these bands were determined and the distribution was calculated as a circular standard deviation (CSD). Lower CSD values represent more aligned – or organized – collagen fascicles; conversely, higher CSD values indicate less organized fascicles. This computational analogue for crossed polarizer imaging has characterized tendon damage in small animal studies [13,15]. Recent work by our group demonstrated that this measurement has strong intra- and interrater reliability in healthy human tendon [*r* > 0.71, accepted manuscript].

Tendon thickness and echogenicity were calculated using established-quantitative methods [6,17,18]. The longitudinal thickness of the tendon was measured as the point to point distance between the deep and superficial edges of the tendon. These measurements were made by a single investigator using an open-source image analysis tool (ImageJ, version 1.51k) [18]. Previous work has shown measurements of longitudinal thickness to be highly reliable between investigators (r > 0.96) [6]. Average echogenicity was calculated for the same tendon regions of interest that was used to calculate CSD, by averaging each pixel grayscale value.

To test our hypothesis that the Achilles tendons of competitive-distance runners differed structurally without presentation of tendinopathic symptoms than their recreationally-active peers, we compared measurements of collagen organization, tendon thickness, and echogenicity between the two groups using one-way unpaired t-tests. The a-priori significance level of 0.05 was adjusted to account for multiple comparisons using a Bonferroni correction (α = 0.05 / 3 = 0.017). Secondary analyses were performed to determine if other runner characteristics, sex, BMI, self-reported foot-strike pattern (forefoot, mid-foot, and rearfoot), and years running at the collegiate level explained measurements of collagen organization. Unpaired two-way t-tests were performed to test differences explained by sex and foot-strike pattern. Univariate linear regression was performed to test the relationship between the continuous variables of BMI and years running with collagen alignment.

## Results

Runners had high VISA-A scores (93.4 ± 8.05, N = 19) and no signs of neovascularization in the tendon (Table 2). Three runners did not complete the VISA-A score and were excluded in further analyses despite meeting our inclusion criteria, which included no current or recent Achilles tendon pain. Some runners had reported dealing with running related pain either at the time of the study or previously; however, these reported events did not originate the Achilles tendon. All runner tendons were qualitatively graded as having ‘normal structure’ and only one of the Doppler scans showed ‘mild neovascularization’, while the remaining subjects in the runner cohort showed no signs of neovascularization.

**Table 2.**
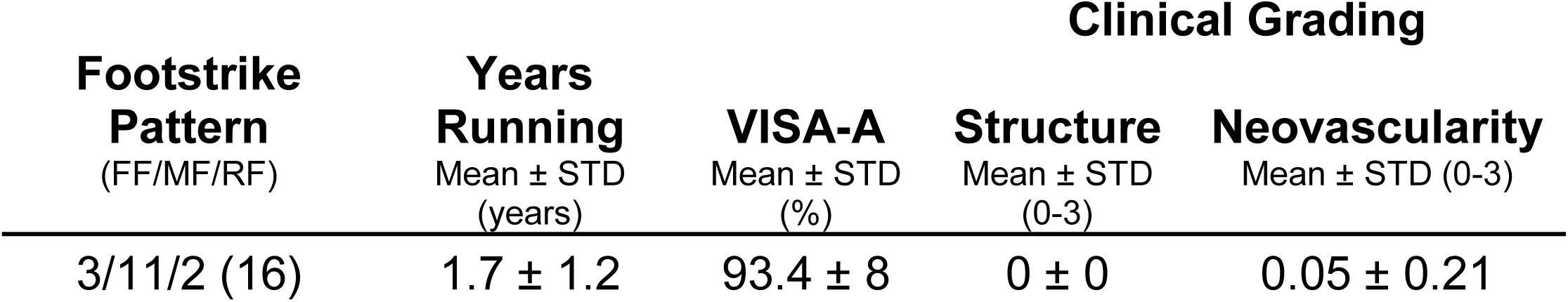
Runner information and tendon clinical grading

Collagen alignment (CSD) was 40% less organized in the runners compared to controls (*p* < 0.001, Fig. 1). Additionally, collagen alignment measures in the runners were 64% less variable, quantified as the coefficient of variation, than those in the controls. Sex, BMI, self-reported foot-strike pattern, and years running at the collegiate level had no effect on measurements of collagen alignment in the competitive runners (*p* > 0.1). Ultrasound images of runner tendon demonstrated darker and thicker tendon mid-substances compared to the control subjects (Fig. 2). Images of the runner tendons were 41% less echogenic (p < 0.001) compared to control scans. Mean echogenicity (Fig. 3) was negatively correlated with collagen alignment in the runner group (R^2^ = 0.24, p = 0.03) but not amongst the control subjects (R^2^ = 0.02, p = 0.68). Measurements of tendon mid-substance thickness were 48% greater in runners than controls (p < 0.001, Fig. 1). However, there were no signs of fusiform swelling of the tendon mid-substance that characterizes tendinopathy.

**Figure 1:**
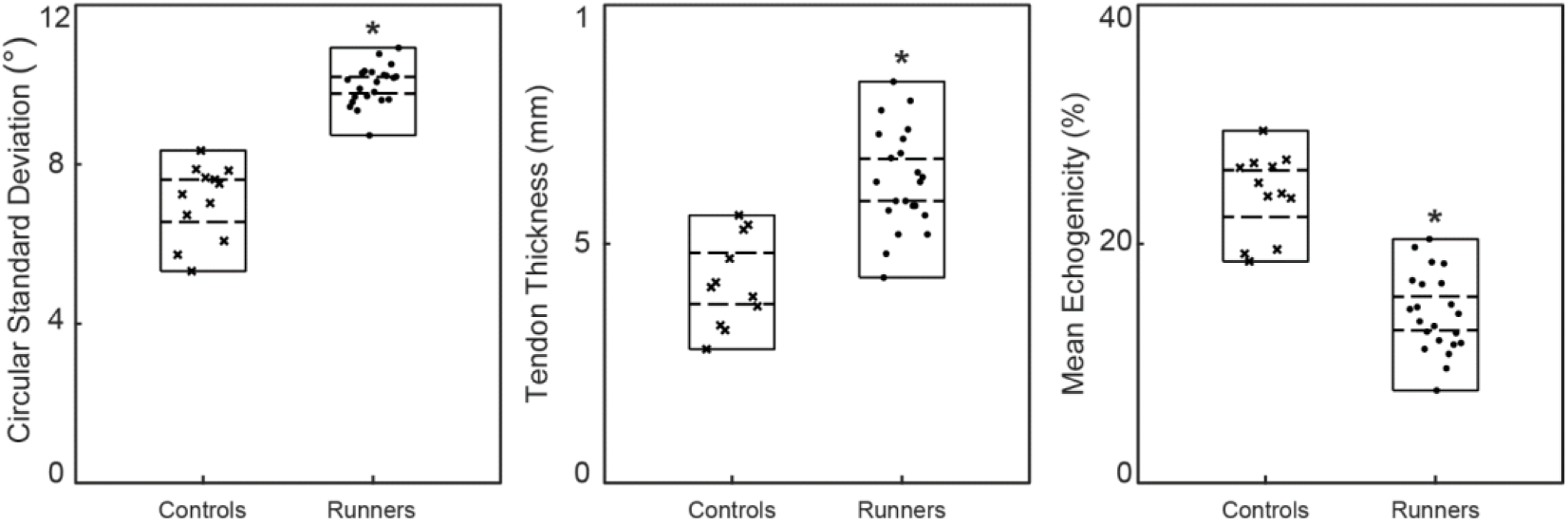
Measures of Circular Standard Deviation (left), Tendon Thickness (center), and Mean Echogenicity (right) are plotted for runners (dots) and controls (crosses). The measurement ranges (solid) and 95% confidence intervals (dashed) for each group are plotted as well. Runners had 40% less organized, 48% thicker, and 41% less echogenic tendons compared to controls (p < 0.001).

**Figure 2:**
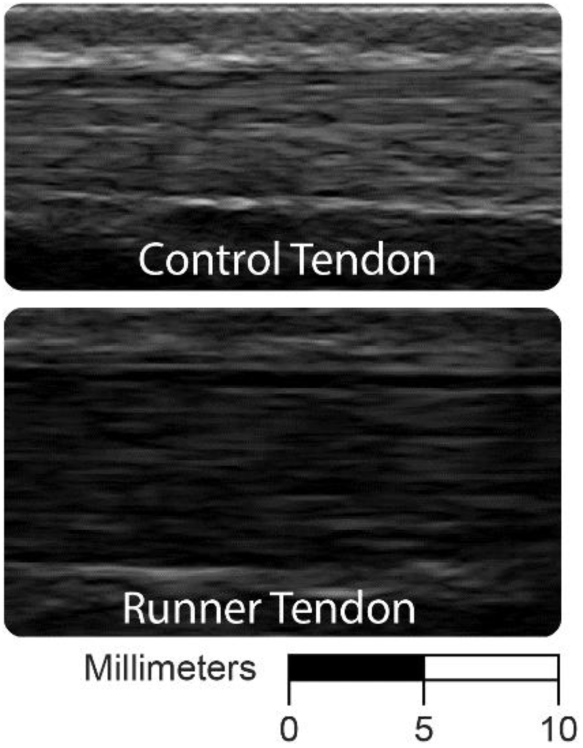
Major differences between the runner and control group can be seen when comparing these tendons under ultrasound. Runner tendon appears visibly thicker and less echogenic than control tendon.

**Figure 3:**
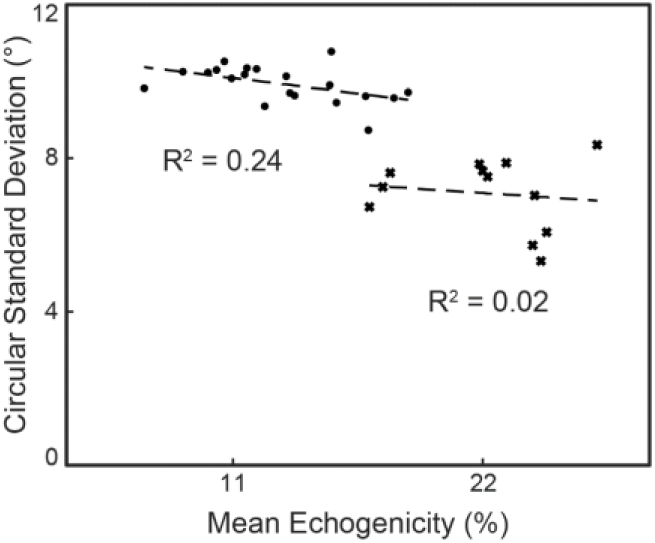
Mean echogenicity was compared to Circular Standard Deviation for the runners (dots) and controls (crosses). There was a negative correlation between ME and CSD for the runners but not for the controls.

## Discussion

Achilles tendon structure differs in competitive-distance runners compared to recreationally-active young adults despite no clinical presentation of mid-substance tendinopathy or pain. Specifically, ultrasound analyses of the Achilles tendon mid-substance highlights reduced collagen alignment and echogenicity and increased tendon thickness in competitive-distance runners. Compromised tendon material properties are partly mitigated by tendon thickening in individuals with Achilles tendinopathy [3], suggesting a compensatory response to maintain tendon strain during high-stress activities. These differences in tendon structure are not explained with risk factors commonly associated with musculoskeletal pain [19] such as sex, BMI, or foot-strike pattern (R^2^ < 0.01). Additionally, less-variable tendon structure in runners compared to controls suggests that these structural changes may be highly dependent on the biomechanical demands placed on tendon, which is similar amongst distance runners on the same team [20].

Differences in Achilles tendon structure observed in this study agree with prior reports of Achilles tendinopathy development and progression and adds new insights into the differences that exist between populations. This cross-sectional study quantified tendon structure of competitive-distance runners at a single time point just prior to the start of a competitive season. Although distance runners are at a 10-fold risk of developing tendinopathy [16], the majority of runners never report symptoms or pain [21]. Therefore, we propose two divergent structural responses to the biomechanical rigors of distance running: habituated tendon and symptomatic tendon. Our results suggest that cyclic tendon loads elicit structural changes in the tendon, which have been linked with decreased tendon properties [17,22]. Tendon hypertrophy appears to mitigate decreases in tendon material properties as large as 51% [3]. However, it remains unclear as to what sustained biomechanical environment results in symptomatic or habituated and pain-free tendons. Training and exercise has been shown to increase tendon thickness [9,10,23], and elevated levels of peritendinous collagen synthesis are responses to short term [24] and long term [25] exercise. Thus, targeted interventions that improve healing responses and beneficial remodeling may mitigate the risks of developing tendinopathy during high-risk activities such as distance running.

Our findings suggest that cyclic tendon loading, experienced during distance running, drives changes in tendon structure without leading to symptoms of tendinopathy. This is highlighted by the low variance in collagen alignment measured in runners (Fig. 1), which may be a normal response to the cyclic loading experienced during distance running. Achilles tendon stiffness also scales with plantarflexor strength [26–28], suggesting that the tendon undergoes remodeling as a protective mechanism against excessive strain in order to maintain good health. In addition, the absence of painful symptoms, neovascularization, and perceived functional deficits in the runners indicates a lack of pathology [6].

Reliable and non-invasive techniques that quantify tendon collagen organization provide opportunities to develop imaging biomarkers that could forecast the development of symptomatic tendinopathy. Currently, tendinopathy is difficult to predict and only detected after symptoms manifest, at which point, treating the condition are not always successful [29–32]. Shear-wave elastography detects changes in the mechanical properties of an affected tendon [33] but requires specialized imaging equipment not available in all clinical settings. Other techniques correlate tendon tension with the speed at which sound travels through tendon, but these measures do not directly assess structural changes [34,35]. In contrast, quantifying collagen alignment using ultrasonography only requires a B-mode ultrasound image that any sonographer or physician could acquire without additional training or specialized equipment available in most sports medicine practices.

There are some key limitations to consider for this study. The competitive-runners were collegiate athletes and therefore did not vary greatly in age, BMI, or years running at the collegiate level. While previous work has shown that these factors are not correlated with the development of tendinopathy [1], the lack of correlation between these factors and the CSD observed in this study could be partially attributed to the low variability. Achilles tendon function was not quantitatively assessed, instead we collected VISA-A scores [36] in the competitive-runners and clinically analyzed ultrasound images [6] to confirm that no underlying tendinopathy was present. A functional task such as a single leg heel-raise or a maximal plantarflexion contraction normalized between subjects could be more sensitive to changes in function than a qualitative questionnaire, but the time constraints of this study prevented more thorough biomechanical assessments. However, asymptomatic tendinopathy has not been linked to losses in plantarflexor function, and good clinical outcomes reported by the runners suggests that these structural changes did not lead to a perceived loss in functional ability.

In conclusion, young adults who participate in competitive-distance running have structurally different tendons than recreationally-active young adults. These differences do not appear to be correlated with age, sex, BMI, self-reported foot-strike pattern, or years running at the collegiate level. It remains unclear if these measurements of decreased tendon organization predict tendinopathy symptoms; however, trained runners are at an 10-times increased risk of Achilles tendon injuries [16,37]. The non-invasive imaging technique used to quantify tendon structure can be used to track changes in tendon structure and better characterize the progression of tendinopathy. Prospectively quantifying tendon structure throughout a training season of distance running may identify imaging biomarkers that can predict painful tendinopathy before symptoms manifest.

## Acknowledgments

We would like to thank Annelise Slater for assistance in data collection. Dr. Casey is now an attending physiatrist at Hospital for Special Surgery.

